# On the capacity of putative plant odorant-binding proteins to bind volatile plant isoprenoids

**DOI:** 10.1101/2021.03.02.433599

**Authors:** Deborah Giordano, Angelo Facchiano, Sabato D’Auria, Francesco Loreto

## Abstract

Plants use odors not only to recruit other organisms for symbioses, but to ‘talk’ to each other. Volatile organic compounds (VOCs) from “emitting” plants inform the “receiving” (listening) plants of impending stresses or simply of their presence. However, the receptors that allow receivers to perceive the volatile cue are elusive. Most likely, plants (as animals) have odorant bind proteins (OBPs), and in fact few OBPs are known to bind “stress-induced” plant VOCs. We investigated whether OBPs may bind volatile constitutive and stress-induced isoprenoids, the most emitted plant VOCs, with well-established roles in plant communication. First, we performed a data base search that generated a list of candidate plant OBPs. Second, we investigated *in silico* the ability of the identified candidate plant OBPs to bind VOCs by molecular simulation experiments. Our results show that monoterpenes can bind the same OBPs that were described to bind other stress-induced VOCs. Whereas, the constitutive hemiterpene isoprene does not bind any investigated OBP and may not have an info-chemical role. We conclude that, as for animal, plant OBPs may bind different VOCs. Despite being generalist and not specialized, plant OBPs may play an important role in allowing plants to eavesdrop messages sent by neighboring plants.

## Introduction

Plants synthesize a variety of volatile organic compounds (VOCs) that are important for reproduction and defense, and in general to communicate with other organisms (Ninkovic et al., 2020). Insects and generalist herbivores, or carnivore insects that are also attracted by the volatile “cry for help” released by plants upon herbivore attacks, are all able to sense plant volatiles (Dicke and Loreto, 2010).

Whether volatiles are also important in plant-plant communication is a more fascinating, yet controversial, issue (Vickers et al., 2009). Growing reports show that volatiles are able to influence plant-plant relationships (Baldwin et al., 2002; Erb, 2019; Ninkovic et al., 2019), and that volatiles elicited in “emitting” plants by abiotic or biotic stresses prime defensive responses in non-elicited “receiving” plants (Zuo et al. 2019; Frank et al. 2021). However, no study has so far looked for the primary events in such elusive plant-plant interaction, i.e. the receptors by which plants may perceive the volatiles emitted from neighboring plants are largely unknown.

Recently, it has been proposed that the passage of VOCs across the plasma membrane relies on their active transport. In particular, the presence of an ABC carrier protein involved in active transport into plant cells has been hypothesized (Adebesin et al., 2017). Plant volatile receptors may belong to a similar category of transporters. An alternative explanation is that plants use OBPs as protein carriers, alike animals. Indeed, there have been at least three cases in which the presence of OBP was postulated in plants. These are: a) the COI1 assembly with a jasmonate Zinc finger Inflorescence Meristem (ZIM)-domain (JAZ) protein family (COI1-JAZ), a high-affinity receptor protein for methyl-jasmonate (MeJa), the volatile moiety of jasmonic acid (JA) (Sheard et al., 2010). After JAZ degradation, transcription factors (TFs) are released, which activate downstream genes and the defensive metabolites in plants challenged by abiotic and biotic stresses (Cheong and Choi, 2003). b) The SA-binding protein 2 (SABP2), an esterase of the a/b-fold hydrolase superfamily, that binds salycilic acid (SA) with high affinity, and then converts the biologically inactive methyl ester of SA (MeSA) to active SA inducing systemic acquired resistance (SAR) in plants challenged by stresses (Park et al., 2008). c) The TOPLESS-like protein (TPL) that specifically binds β-caryophyllene, a stress-induced sesquiterpene and a volatile signal for herbivores and carnivores in multitrophic interactions. TPL and TPL-related (TPR) proteins are transcriptional co-repressors (also toward JA-mediated signaling). Interestingly, only the capacity to bind β-caryophyllene was tested with emitting and receiving (eavesdropping) plants (Nagashima et al., 2019).

These three cases need confirmation, and all other plant volatiles (at least 1700 known so far, Dicke and Loreto (2010)) wait for receptor recognition (if any). We report here an *in-silico* study based on current knowledge of plant protein structure, especially aiming at selecting best candidates as plant OBPs for plant volatiles whose receptors are still unknown/unavailable. We particularly focused on the volatile isoprenoids (isoprene and monoterpenes) produced by the methyl erythritol phosphate (MEP) chloroplast pathway and representing the largest component of plant VOC emissions in the atmosphere (Loreto and Schnitzler, 2010).

## Results and Discussion

We looked at plant OBPs following a two-level structural approach (identification of candidate plant OBPs, and validation by *in silico* molecular simulations of OBP capacity), as detailed in the Method section below (see also Figure 1). For the first level:

**Figure 1.**
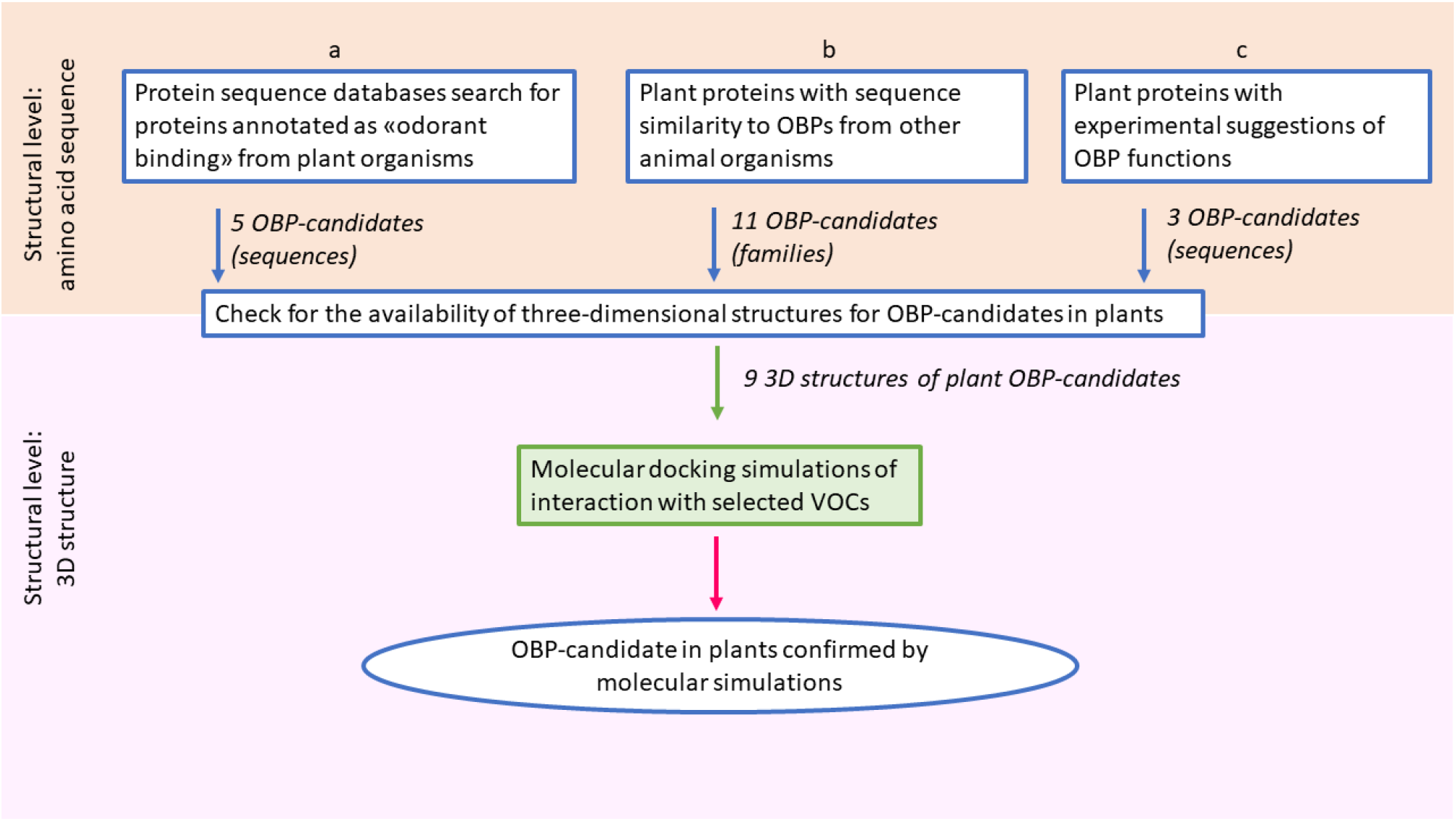
Schematic workflow for the search of candidate odorant-binding proteins in plants. The search was performed at two levels of investigation. The first level was the identification of candidate plant OBPs, based on features and similarities of protein sequences from public data bases (boxes labelled a and b) and on literature evidences (box labelled c). This level identified proteins or protein families, for which amino acid sequences are available. The second level was the validation by *in silico* molecular simulations, to verify the potential capability of the candidate proteins to bind selected VOCs. This level can be applied only to the candidate proteins, selected from the first level, for which the three-dimensional structure is made available by experimental studies.

*First,* a search for plant proteins potentially involved in plant VOC binding was performed in protein sequence databases. Five complete or partial protein sequences were found, three from *Anthurium amnicola* (named OBP56d, OBP A10_1, and OBP A10_2), one from *Nymphaea thermarum* (named putative OBP) and one from *Pyrus x bretschneideri* (named OBP-70 like) (Supplementary Table S1).

However, when these sequences were used for a further screening for similarities with annotated plant proteins in the sequence databases, a statistically relevant similarity was only found in the case of the putative OBP from *Nymphaea thermarum.* In this case the Flowering locus T (FT) and T1 proteins, and the heading date 3A and 3B showed high similarity and coverage with the putative OBP.

By comparing plant and animal OBPs, similarities were identified between OBPs from insects and plant OBP56d and OBP-70 like, between chemosensor proteins and plant OBP A10_1, and between phosphatidylethanolamine-binding proteins and the putative OBP from *Nymphaea thermarum* (Supplementary Table S2).

*Second,* we compared 432 OBP protein sequences from different animal sources with plant protein sequences. Only five animal protein families share sequence similarity with plant proteins, as summarized in Table 1.

**Table 1.** Similarities of animal and plant protein families with odorant receptor/transporter/channel functions.

| Animal protein family | Similar plant proteins |
| --- | --- |
| Bovine Cyclin Nucleotide Gating Olfactory Channel of *Bos Taurus* | Potassium Voltage-Gated Channel |
| Chloride channel Anoctamin-2 of *Mus musculus* | anoctamin-like protein |
| Human/Mouse Receptor Expression Enhancing molecules | HVA22 e HVA22-like plants proteins |
| BPI fold containing family B member 3 from *Mus musculus* and *Rattus norvegicus* | putative BPI, lipid-binding protein, hypothetical protein, and unnamed protein |
| putative OBP 5a from *Drosophila melanogaster* | FT, D3-like, protein “Mother of FT and TFL1-like (Terminal flower 1-like)” (MFT), ZCN9 (MFT-like), Heading Date 3A-like |

It is interesting to note that all plant protein sequences reported in Table 1 are related to inflorescence signaling. HVA22 is induced by ABA/stress and has a role in the GA-induced cellular death and in the regulation of seed germination (Shen et al. 2001). FT is a florigen that induces and promotes the transition from vegetative growth to flowering (Koornneef et al., 1998). The protein MFT is involved in regulation of seeds germination by abscisic acid (ABA)/gibberellic acid (GA) signaling (Vaistij et al., 2018). Heading date 3A-like, as FT, is a probable florigen, which promotes the transition from vegetative growth to flowering downstream of HD1 and EHD1 under short day conditions (Taoka et al. 2011).

The first level of our study, based on comparison of protein sequences, returned a list of plant proteins/families with potential OBP function, to which the three plant OBPs already described (see Introduction) were added. Experimental 3D structures are available for nine of the identified putative plant OBPs. These OBPs were therefore selected for the second level of the study, involving molecular docking simulations of the interactions between potential plant OBPs and selected VOCs, to finally identify candidate plant OBPs.

Table 2 reports the binding energy values obtained by docking simulations for each complex between potential plant OBPs and ligands (plant VOCs), together with the binding energy values obtained as a reference for experimental complexes.

**Table 2.** Binding energy values, in Kcal/mol, obtained by docking simulations, between putative plant OBPs and isoprenoid VOCs.

| OBPs | α-Pinene | Limonene | Myrcene | β-Caryophyllene | Isoprene | Linalool |
| --- | --- | --- | --- | --- | --- | --- |
| TPL-like | -5.06 | -4.76 | -3.95 | -6.16 | -3.39 | -3.94 |
| ABA receptor | -4.69 | -4.73 | -3.84 | -6.32 | -3.35 | -4.12 |
| GA receptor | -5.50 | -5.44 | -4.92 | -6.94 | -3.78 | -4.69 |
| Heading Date 3A | -5.46 | -4.99 | -4.29 | -6.83 | -3.44 | -4.79 |
| FT | -5.23 | -5.07 | -4.26 | -6.65 | -3.15 | -4.98 |
| TFL1 | -5.30 | -5.53 | -4.38 | -7.15 | -3.70 | -4.98 |
| Partial JA receptor | -5.23 | -4.79 | -4.16 | -6.05 | -2.78 | -4.21 |
| Complete JA receptor | -5.92 | -5.11 | -4.41 | -6.61 | -2.82 | -4.49 |
| SABP2 | -6.03 | -6.12 | -5.14 | -6.73 | -3.25 | -4.70 |
| Reference protein * | -5.42 | -6.29 | -4.15 | not available | not available | not available |

\* Reference protein is the protein for which it has been found an experimental complex with the ligand. For α-pinene, limonene, and β-myrcene, the reference proteins are cytochrome P450 2B6 complexed with α-pinene (PDB code: 4I91), lipid binding protein complexed with limonene 1,2 epoxide (PDB code: 2A2G, and linalool dehydratase/isomerase complexed with β-myrcene (PDB code: 5HSS), respectively.

The results suggest that the three monoterpenes tested (α-pinene, β-myrcene and limonene) may bind some of the putative OBPs with energy values similar or lower than the values observed for the reference complexes. In fact, α-pinene binds the reference protein cytochrome P450 2B6 with an energy value of −5.42 Kcal/mol, and a predicted Ki of 103 μM (predicted Ki values are in Supplementary Table S3). In our docking simulations, the best results for α-pinene were obtained with SABP2 and the complete JA receptor, with binding energy value of −6.03 Kcal/mol and −5.92 Kcal/mol, and predicted Ki of 37 μM and 38,55 μM, respectively. In both cases, this is a better interaction than with the reference complex. In the case of GA receptor and heading date 3A, α-pinene has binding energy values very similar to the value in the reference complex.

Similarly to α-pinene, β-myrcene binds SABP2, GA receptor and the JA receptor better than the reference complex. Among the other candidate proteins, protein heading date 3A, FT, and tfl1 showed binding energy values similar to the reference complex for β-myrcene.

In the case of the other reference complex, the ligand reported in the literature (limonene 1,2 epoxide) is a modified form of the plant volatile used in our simulation experiments (the monoterpene limonene). Therefore, we used this reference complex with less confidence. In any case, the energy values were similar to the reference complex only for SAPB2.

Results obtained for β-caryophyllene, isoprene, and linalool could not be compared to a reference complex. The binding energy values might be different for the three compounds due to their chemical and structural properties. However, remarkably β-caryophyllene showed the lowest, and isoprene the highest, energy binding values.

Our analysis overall confirms that OBPs might be present in plants and may also bind VOCs produced by plants through the methyl-erythritol-phosphate (MEP) pathway. MEP synthesizes the volatiles isoprene and monoterpenes, both constitutively and after induction by stress agents (Dicke and Loreto, 2010). While monoterpenes are efficiently bound by OBPs, isoprene, the simplest and most abundantly emitted volatile isoprenoid, does not seem to bind efficiently any OBP. This is in line with ecological observations reporting a role for monoterpenes in plant communication with other organisms (Bouwmeester et al., 2019) which is arguably not observed for isoprene (e.g. Brilli et al., 2009), However, isoprene influences many plant traits (recently reviewed by Monson et al., 2020). Despite absence of OBP binding capacity, isoprene profoundly modifies properties of cellular and sub-cellular membranes (Velikova et al. 2016, Pollastri et al. 2019) which may in turn activate signals reshaping plant genomes and phenomes (Harvey and Sharkey 2016; Miloradovic van Doorn et al. 2020).

Interestingly, monoterpenes seem to bind more efficiently with OBPs that are also reported to bind other plant volatiles. In particular, SABP2, the SA-binding protein that strongly binds the stress-induced volatile MeSA, also seems to be a candidate for the three monoterpenes tested. Protein heading date 3A and tfl1, GA receptor and FT may also bind, perhaps more specifically, the three monoterpenes. However, our results indicate that, as reported for the OBPs from animals and insects, the candidate plant OBPs have a broad ligand binding specificity, and, consequently, they may bind several different VOCs (Ramoni et al., 2007).

In many cases, binding of the ligands occurs at the same protein structure site, as shown for SABP2 in the experimentally reported complex with SA (Figure 2), and the simulated complexes with α-pinene (Figure 3A) as well as with other VOCs (Supplementary Figures S1 and S2).

**Figure 2.**
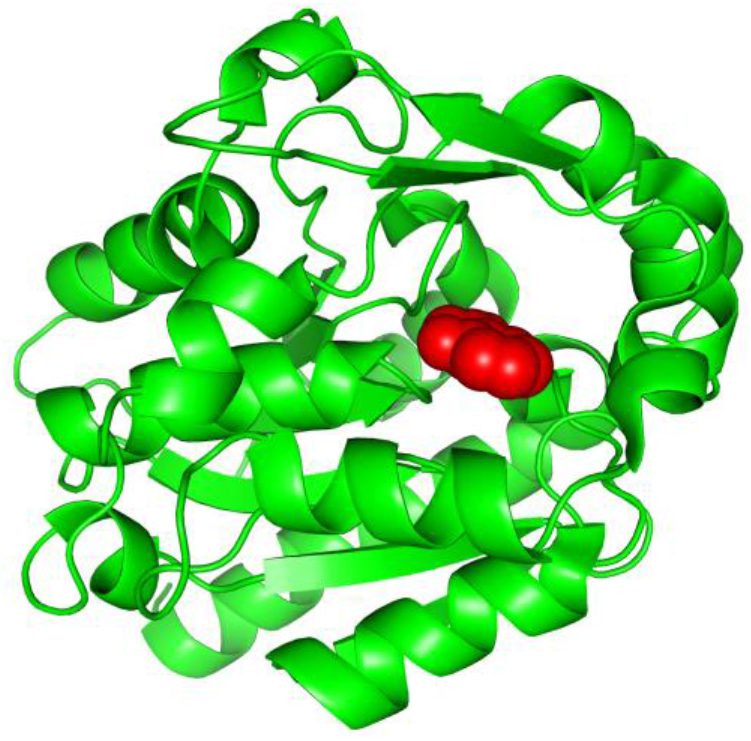
Experimental structure of SABP2 protein structure with salicylic acid in the binding site (PDB structure 1Y7I). The architecture of SABP2 is schematized by the backbone structure (green), with ribbons and arrows to evidence helices and beta strands, respectively. The salicylic acid molecule (colored in red) is in space-fill representation.

**Figure 3.**
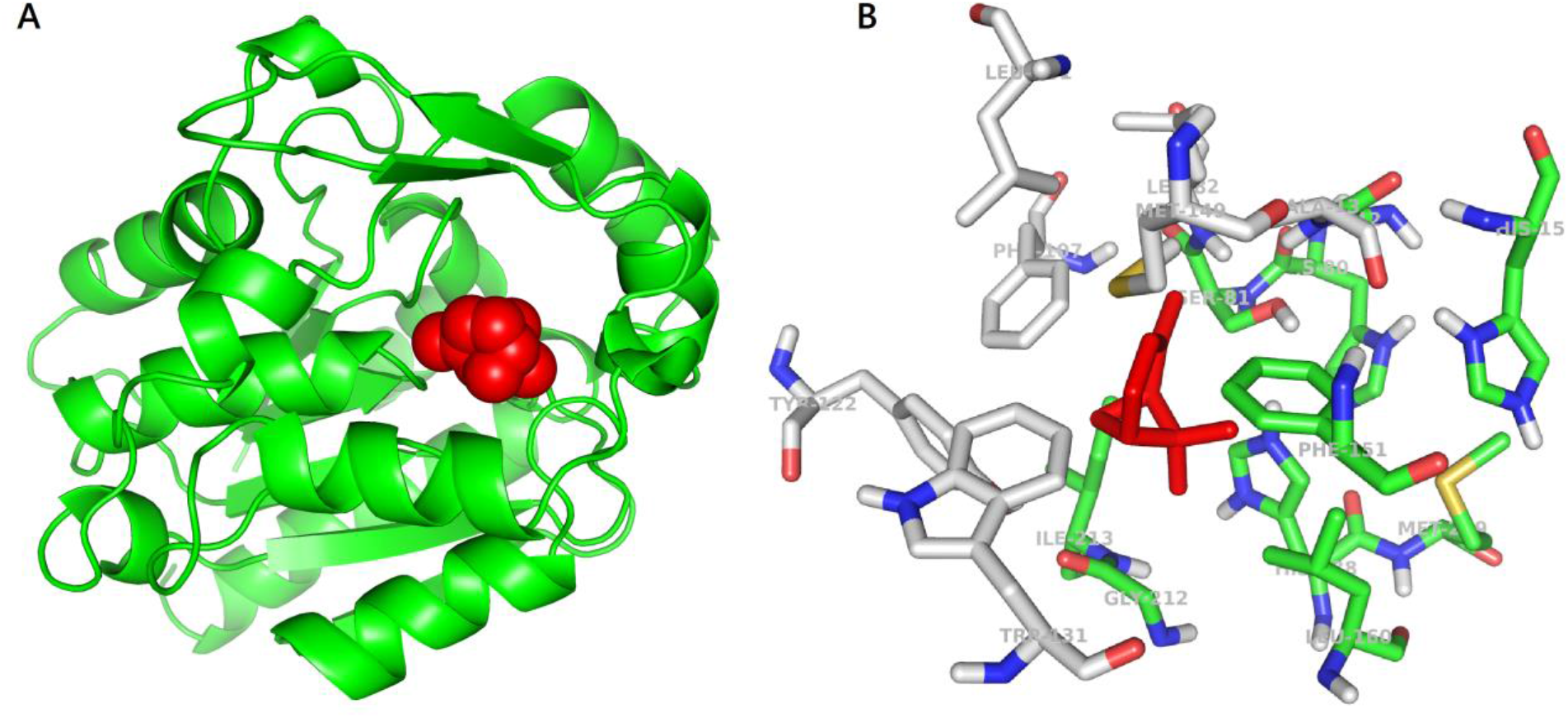
***Panel A:*** Molecular docking simulation of SABP2 protein structure with α-pinene (red molecule) in the binding site. SABP2 is shown with the same spatial orientation of Figure 2 to emphasize that α-pinene occupies the same binding site of salicylic acid. ***Panel B:*** Focus on the binding site of SABP2. α-Pinene (red molecule) interacts directly with amino acid residues labelled with carbon atoms in green. Amino acid residues with carbon atom in grey are also part of the binding site, although not directly in contact with α-pinene. Standard colors are used for the other amino acid atoms (red=oxygen, blue=nitrogen, yellow=sulphur, white=hydrogen).

The SABP2 binding site (Figure 3B) is characterized by the presence of aromatic side chains, also observed in other candidate plant OBPs. In detail, the binding site of SABP2 includes two phenylalanines, one tyrosine, and one tryptophan, while in the GA receptor there are two phenylalanines and four tyrosines, in the JA receptor one phenylalanine, one tyrosine, and one tryptophan residue, and in the FT receptor two phenylalanines and one tryptophan. Other candidate plant OBPs have some aromatic side chains in the binding site, although in lower number (for example, the ABA receptor has one phenylalanine and one tyrosine, while TLF1 has two phenylalanines). This is also reminiscent of the binding site of OBPs from animal organisms (Bianchet et al., 1996). Bovine OBPs includes five phenylalanines and one tyrosine; drosophila OBPs have four phenylalanines and one tryptophan, and porcine OBPs have two phenylalanines. The meaning of this conserved feature across biomes may be related to a common necessity to create a hydrophobic environment where the hydrophobic odorant molecules can be accommodated.

Overall, our study confirms that plant OBPs may exist, and that they may be structurally and functionally similar to OBPs described in animals. As in the case of animal OBPs, also plant OBPs seem to be able to bind different VOCs in the same binding site, using the same amino acid sequences. Retrieval and description of plant OBPs may be an important step to unveil how plants eavesdrop messages sent by other plants and how the information is then used to activate molecular and metabolic changes.

## Methods

The search for potential OBP proteins in plants was performed following the procedure schematized in Figure 1, and consisted of two levels of investigation.

The first level was about amino acid sequences to search for plant proteins with potential OBP function. The second level was about experimental 3D structures of the candidate plant OBPs, to validate by molecular simulations their potential ability to bind volatile molecules.

In detail for the first level, three steps (named a, b, and c, see Figure 1) were followed. Step “a” was a screening for proteins of interest performed on the UniProt (http://www.uniprot.org) and NCBI (http://www.ncbi.nih.nlm.gov) protein databases. Initial screening was performed by using the protein name and entry annotations, with the query “odorant binding protein”. Five plant proteins were found, annotated as “predicted proteins”, which means that they were obtained by nucleotide sequence translation, without evidence at protein or transcript levels, and the name was assigned to the proteins by similarity to other proteins. In the case of the Uniprot Entry A0A1D1ZDX5, named general odorant-binding protein 56d (OBP56d), from *Anthurium amnicola,* the annotations revealed that the protein is included into the “Pheromone/general odorant-binding protein superfamily” of the InterPro database (http://www.ebi.ac.uk/interpro/). In the case of UniProt entry A0A1D1Z329, named putative odorant-binding protein A10_1 (OBPA10_1), from *Anthurium amnicola,* the annotations revealed that the protein is included into the “Insect odorant-binding protein A10” protein family of the InterPro database. These observations may explain the OBP annotation for these proteins.

The protein sequence selected were further investigated by BLAST searches for similar sequences, by using the BLAST interfaces at the database web sites. Standard BLAST search parameters were used, which means that results with E-value <10 are reported in the output. This setting leaves a wide possibility of including results not significant, being the E-value < 0.001 considered the reference threshold for similarity.

Step “b” was of a search for plant proteins similar to the 432 OBPs from animal sources available in the protein databases. BLAST searches performed for the 432 OBPs, identified plant proteins and protein families with similarity to known OBPs. Step “c” was about collecting information for plant proteins whose OBP function has been experimentally tested (see Introduction).

The second level of investigation concerned the study by molecular simulations of the interaction of the potential plant OBPs with selected VOCs. We verified the availability of three-dimensional (3D) structures of the candidate OBPs in plants. In particular, the Protein Data Bank (PDB) (www.rcsb.org), collecting the 3D structure of proteins, was interrogated for appropriate protein structures of the candidate OBP from plants resulting from our first level search. The screening allowed us to select the following plant proteins with potential primary or secondary function as OBP, and with available 3D structures: ABA Receptor from *Arabidopsis thaliana* (PDB code: 4dsb), GA receptor GID1 from *Oryza sativa* (3ebl), Flowering locus t (FT) from *Arabidopsis thaliana* (1wkp), Terminal flower 1 (tfl1) from *Arabidopsis thaliana* (1wko), Protein Heading date DATE 3A from *Oryza sativa* (3axy), TPL-like protein from *Arabidopsis thaliana* (5nqs), COI (partial JA receptor) from *Arabidopsis thaliana* (3ogl), COI & JAZ (JA receptor complete) from *Arabidopsis thaliana* (3ogl), salicylic acid binding protein 2 (SA enzyme) from *Nicotiana tabacum* (1y7i)

Molecular structures of small molecules were extracted from the PubChem database. The VOCs selected as ligands in our study were the isoprenoids α-pinene, limonene, β-myrcene, β- caryophyllene, isoprene, linalool.

Molecular simulation experiments of protein-ligand interactions were carried out with Autodock 4.2 and AutoDock Tools 1.5.6 (Morris, 2009), which allowed us to prepare the screening, perform the docking simulation, and analyze the results. The binding energy values obtained for the simulated protein-ligand complexes were compared to the values for complexes used as reference. We found in the PDB database complexes of animal proteins with α-pinene, limonene 1,2 epoxide, and β- myrcene. α-Pinene and β-myrcene are two of the selected VOCs for our simulation, as they correspond to the natural molecules synthesized and emitted by plants. Limonene 1,2 epoxide is a modified form of the natural VOC. Although not identical to the corresponding plant VOC, it may be useful as additional reference value. For available plant receptors (ABA receptor, GA receptor, JA receptor, and SA binding protein 2), the reference structures are complexes with ABA, GA, JA-isoleucine, and SA, respectively. Also, these complexes may offer additional reference values of binding energy.

To validate the docking simulation experimental protocol, we applied a re-docking procedure to the reference complexes, following the procedure in use in our laboratory (Scafuri et al., 2016, 2020). We depleted the ligand from the complex ligand-protein and then the ligand-depleted complex (the protein alone) was used to simulate the ligand docking. The re-docking experiments were carried out for the protein-ligand reference structures selected above. This approach allowed us to check that the simulation procedure located correctly the ligand in the expected binding site, and gave us the reference value of the binding energy expected in the true protein-ligand complex.

Molecular visualization of results was obtained with PyMOL Molecular Graphics System, Version 1.3 Schrödinger, LLC.

## Acknowledgements

FL acknowledges contribution from the Project PRIN - COFIN 2017 (Italian Ministry of University and Research): “ Plant multitROphic interactions for bioinspired Strategies of PEst ConTrol (PROSPECT)”. We would like to thank Prof. Paolo Pelosi for the inspiring discussions about OBPs.

## Supplementary materials

**Supplementary Table S1.**
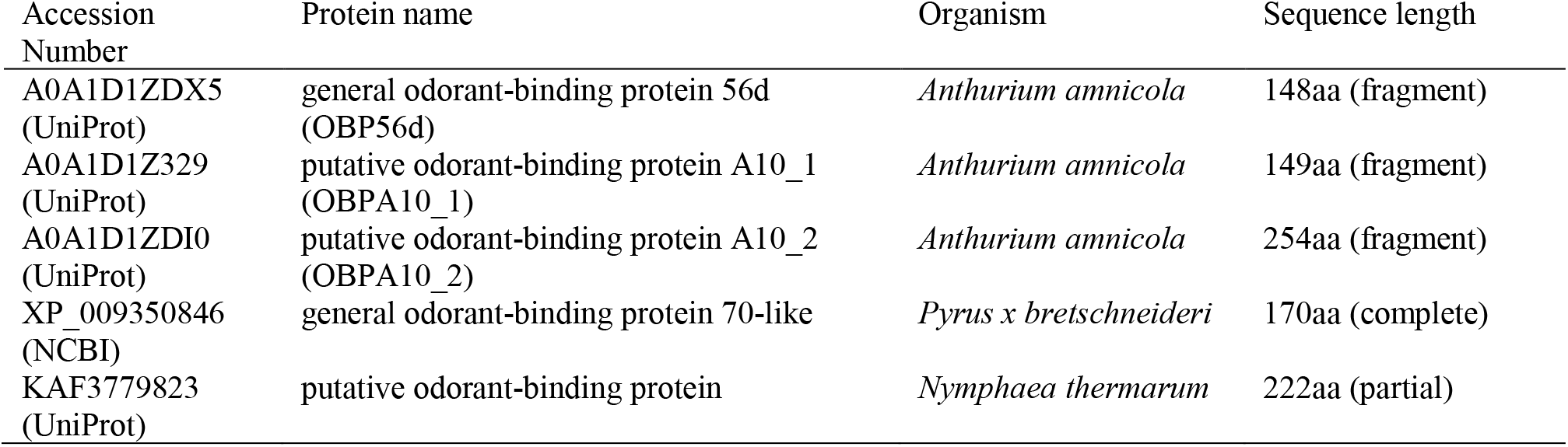
Plant proteins from sequence databases annotated as “odorantbinding protein”.

**Supplementary Table S2.** BLAST results for non-plant proteins with similarity to “general odorant binding protein 56d” from *Anthurium amnicola*. Only the best ten results are shown.

| Accession Number | Annotation | Organism | Identity % | Alignment length | E-Value<sup>a</sup> | Score<sup>a</sup> |
| --- | --- | --- | --- | --- | --- | --- |
| XP_034232292.1 | general odorant-binding protein 56d-like isoform X1 (151 aa) | Thrips palmi | 63.87 | 119 | 4e-51 | 169 |
| XP_026281309.1 | uncharacterized protein LOC113208505 (148 aa) | Frankliniella occidentalis | 50.00 | 128 | 5e-28 | 110 |
| KAE8741848.1 | hypothetical protein FOCC_FOCC012596 (190 aa) | Frankliniella occidentalis | 50.39 | 127 | 2e-27 | 110 |
| XP_034249409.1 | uncharacterized protein LOC117650253 (149 aa) | Thrips palmi | 31.09 | 119 | 3e-11 | 68.2 |
| XP_015835846.1 | PREDICTED: pheromone-binding protein-related protein 6-like (145 aa) | Tribolium castaneum | 30.50 | 141 | 3e-10 | 65.1 |
| EFA04593.1 | odorant binding protein 07 (136 aa) | Tribolium castaneum | 29.85 | 134 | 7e-10 | 64.3 |
| XP_019697001.2 | general odorant-binding protein 83a (132 aa) | Harpegnathos saltator | 25.37 | 134 | 1e-09 | 63.5 |
| KAF2881145.1 | hypothetical protein ILUMI_25019 (123aa) | Ignelater luminosus | 32.23 | 121 | 2e-08 | 60.1 |
| XP_011346593.1 | general odorant-binding protein 83a isoform X1 (133 aa) | Ooceraea biroi | 28.57 | 126 | 3e-08 | 59.7 |
| XP_011164418.1 | uncharacterized protein LOC105199157 (131 aa) | Solenopsis invicta | 29.01 | 131 | 4e-08 | 59.3 |

<sup>a</sup> E-value and score are BLAST measures of significance of the sequence similarity observed. Low E-values (< 0.001) and high score indicate very significant results.

**Supplementary Table S3.**
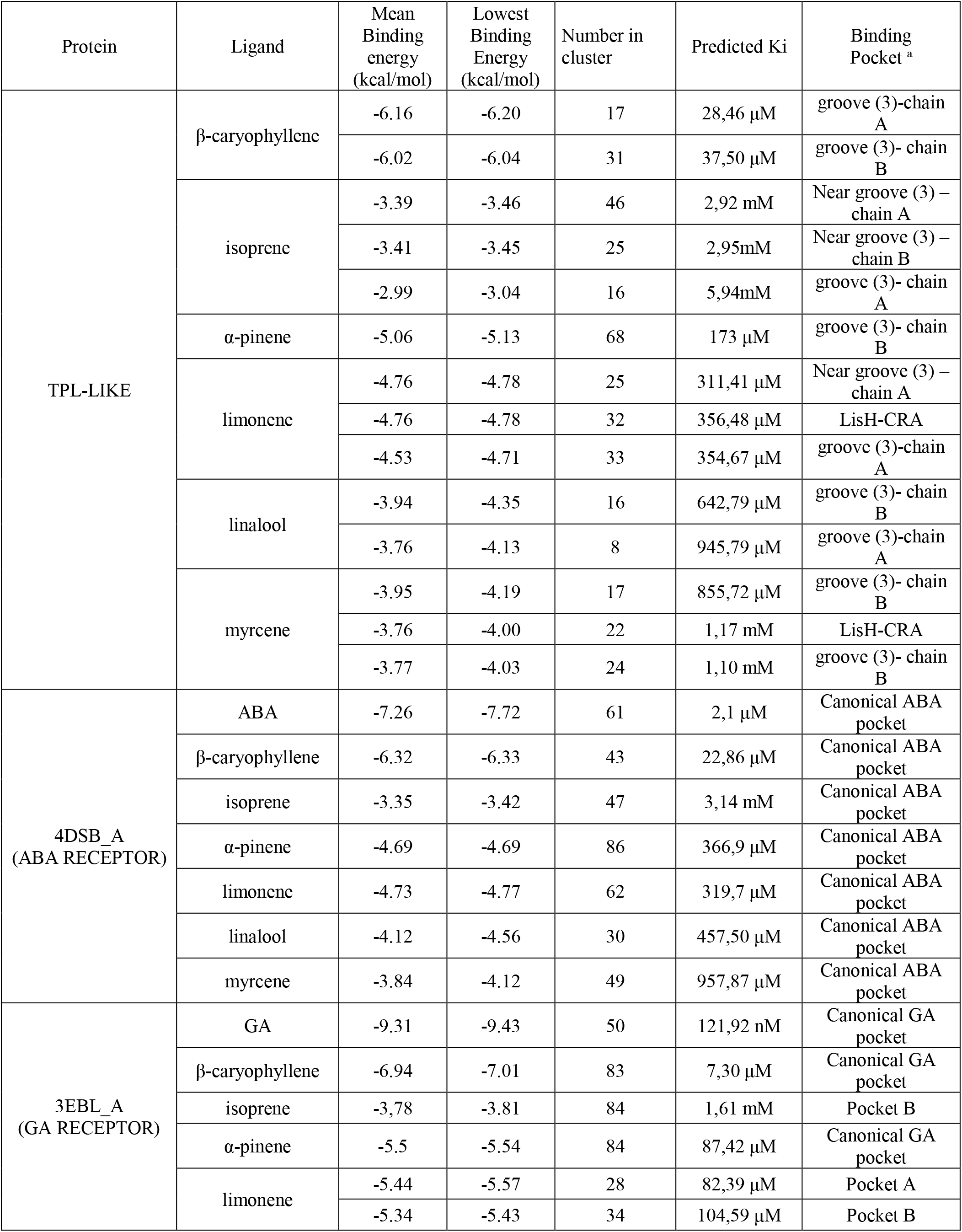
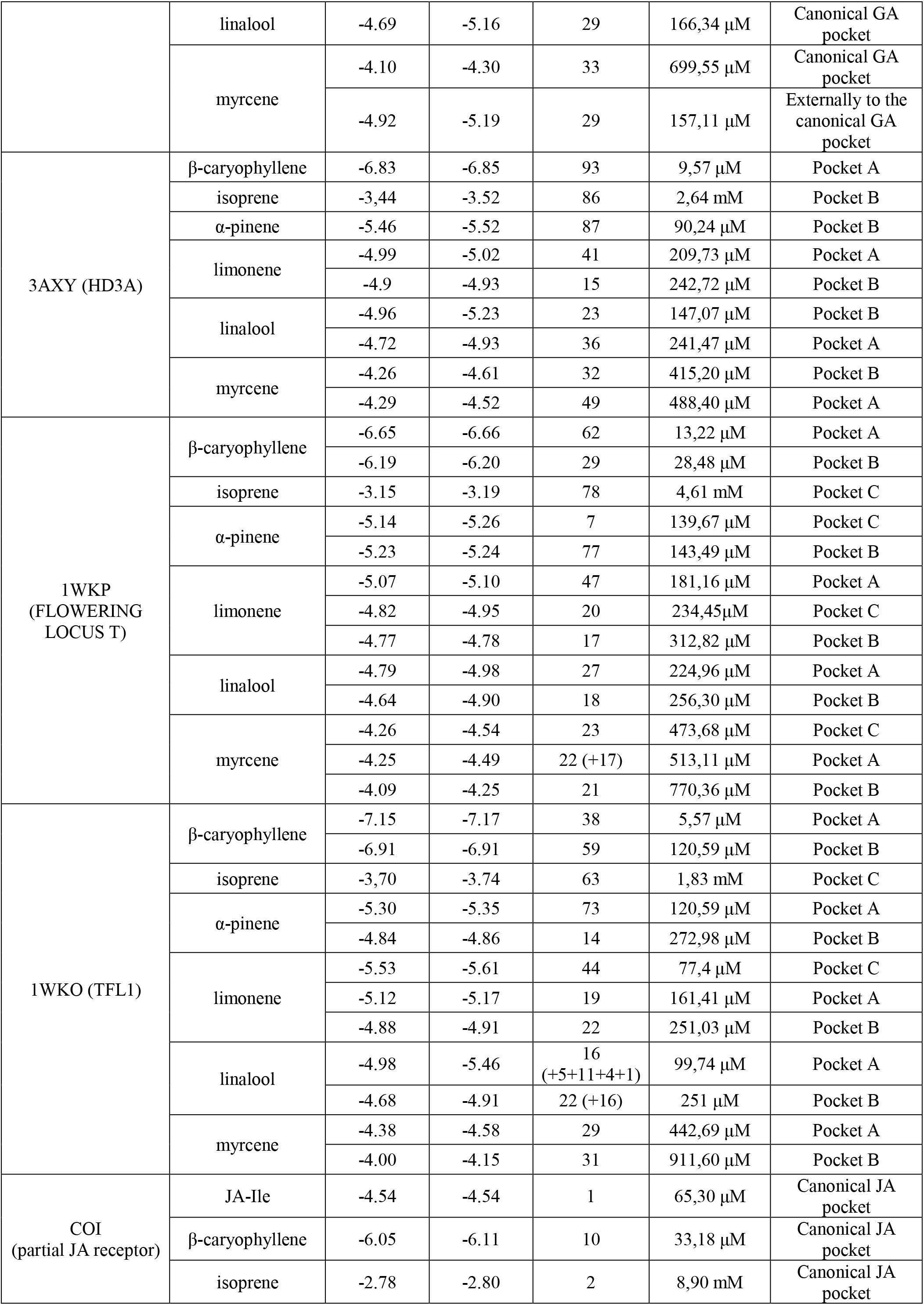
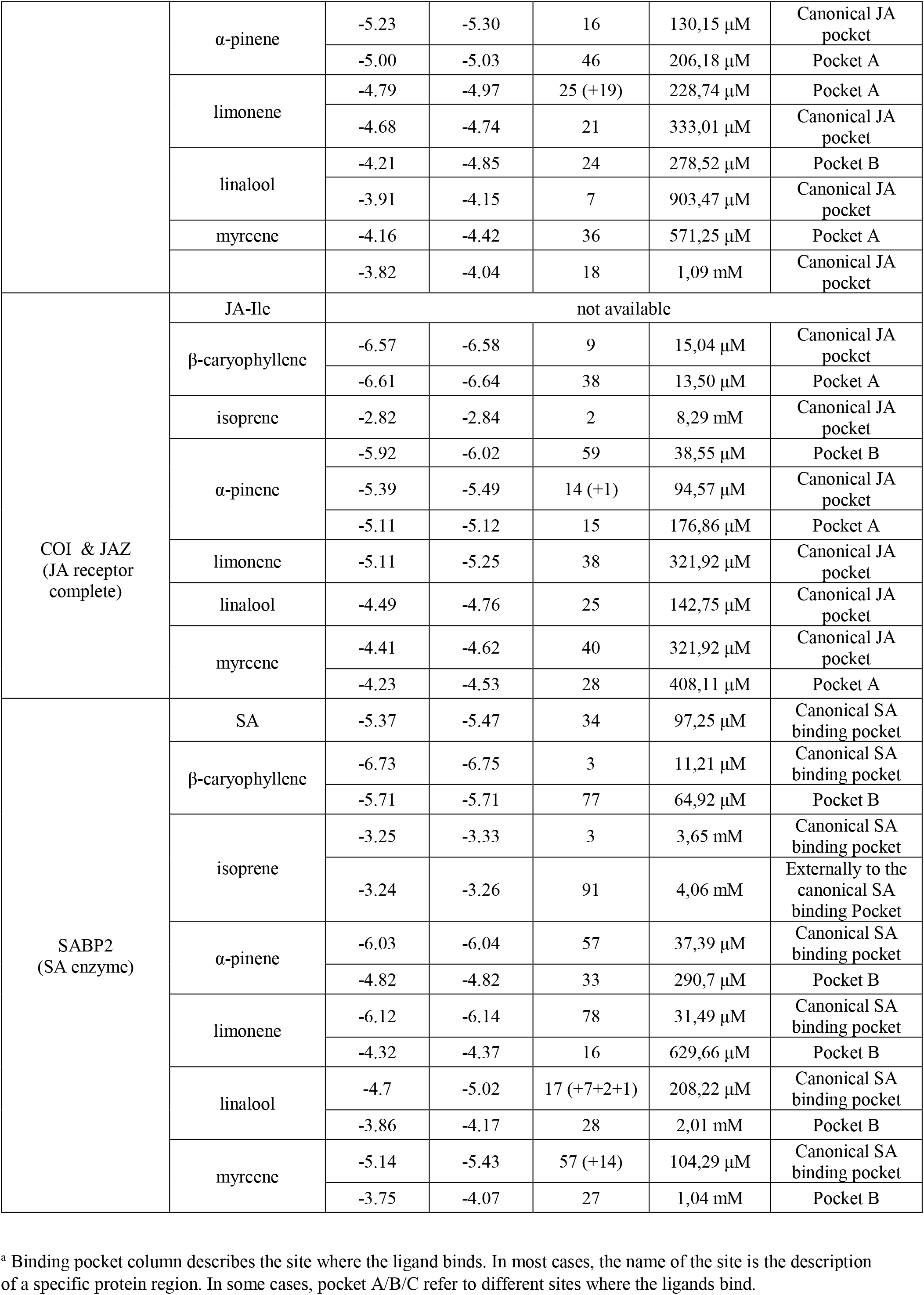
Complete results of docking simulation experiments. When more binding energy and predicted Ki values are reported, they refer to alternative binding sites detected.

**Supplementary Figure S1.**
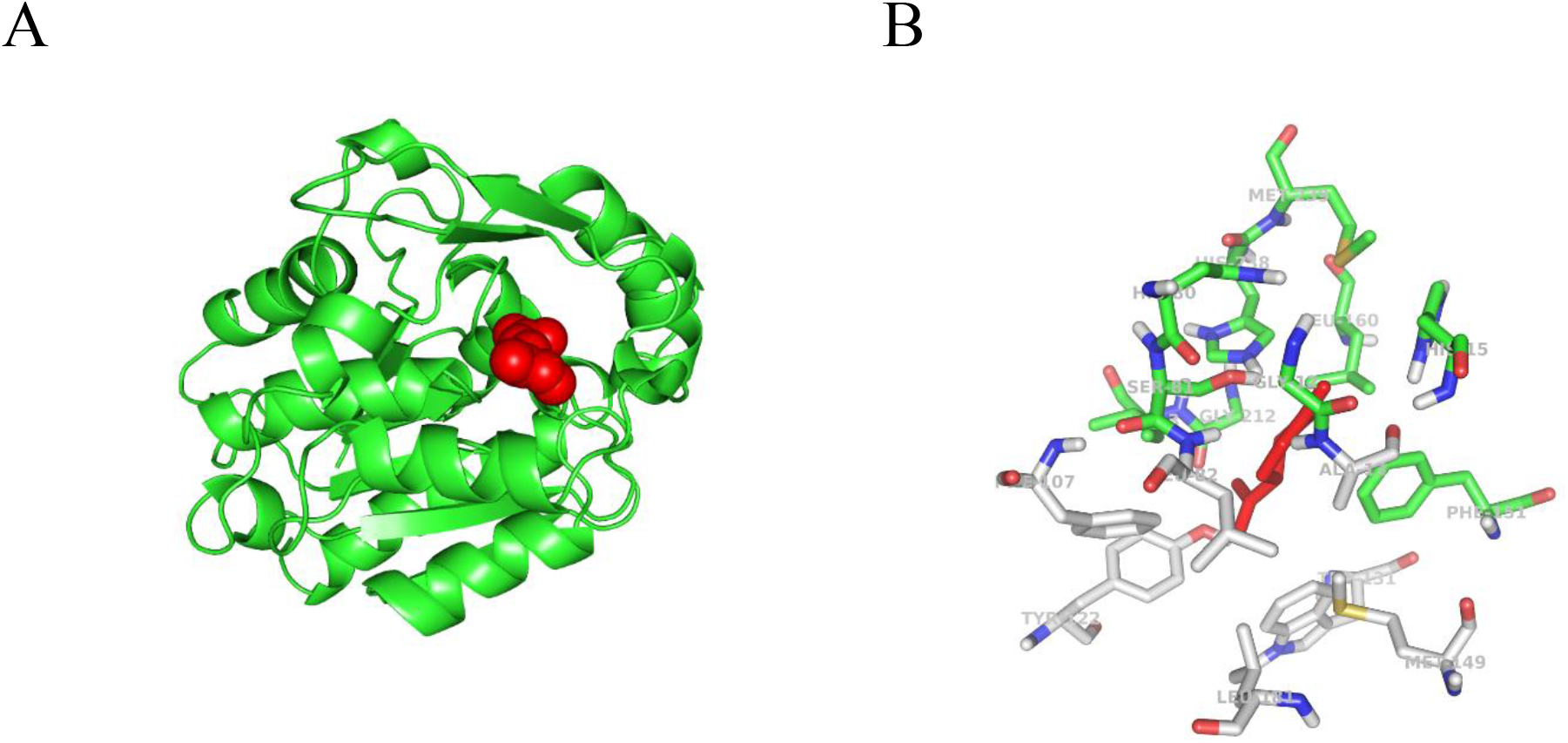
Panel A: SABP2 protein with limonene (red molecule) in the binding site. SABP2 is shown with the same spatial orientation of Figure 2, to evidence that limonene occupies the same binding site of the salicylic acid. Panel B: Focus of the binding site of SABP2. Limonene (red molecule) interacts directly with amino acids labelled with carbon atoms in green. Amino acids with carbon atom in grey are also part of the binding site, although not directly in contact with limonene. Standard colors are used for the other amino acid atoms (red=oxygen, blue=nitrogen, yellow=sulphur, white=hydrogen).

**Supplementary Figure S2.**
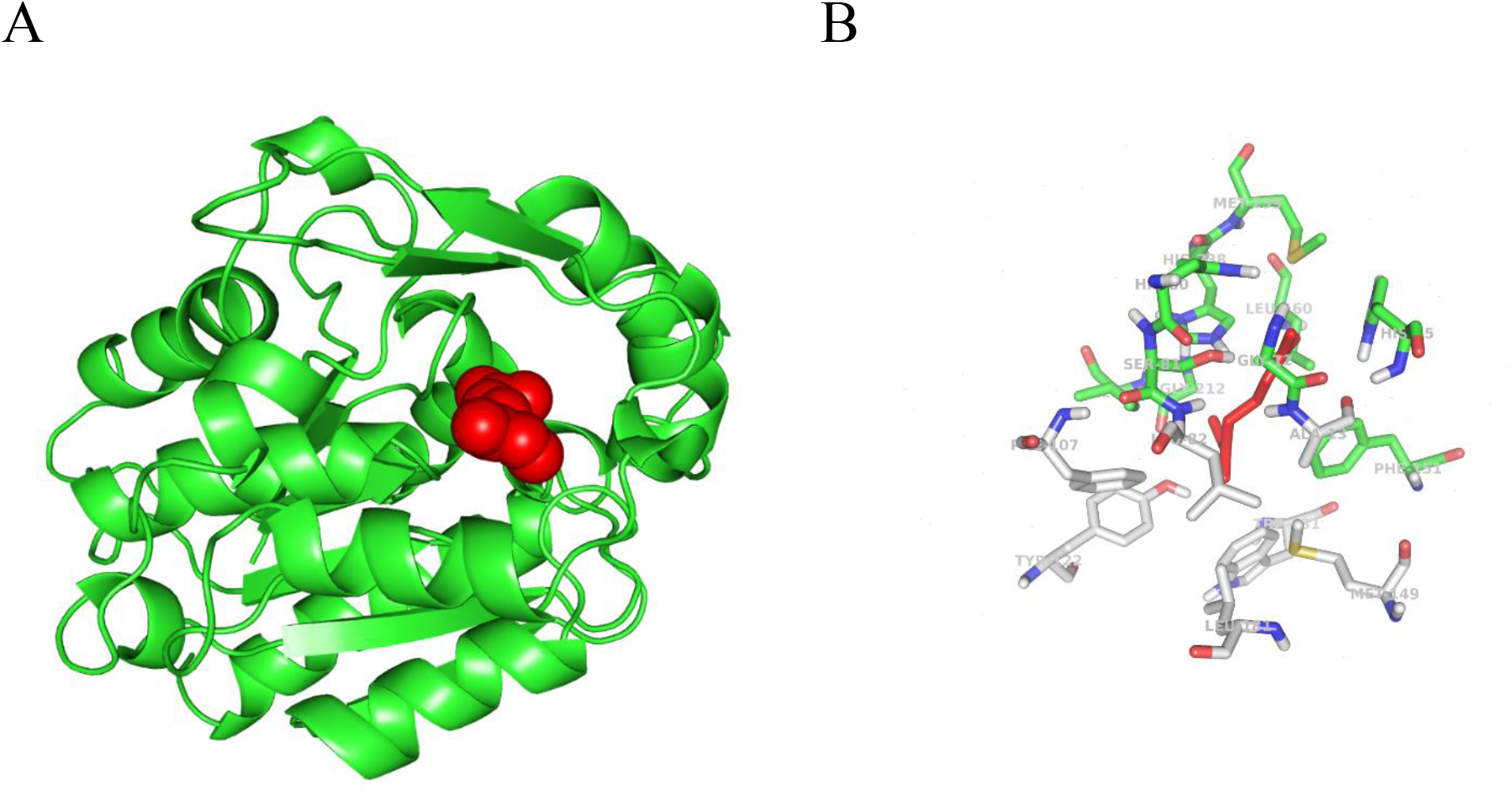
Panel A: SABP2 protein with myrcene (red molecule) in the binding site. SABP2 is shown with the same spatial orientation of Figure 2, to evidence that myrcene occupies the same binding site of the salicylic acid. Panel B: Focus of the binding site of SABP2. Myrcene (red molecule) interacts directly with amino acids labelled with carbon atoms in green. Amino acids with carbon atom in grey are also part of the binding site, although not directly in contact with myrcene. Standard colors are used for the other amino acid atoms (red=oxygen, blue=nitrogen, yellow=sulphur, white=hydrogen).

